# T cells use distinct topological and membrane receptor scanning strategies that individually coalesce during receptor recognition

**DOI:** 10.1101/2022.02.23.481517

**Authors:** En Cai, Casey Beppler, John Eichorst, Kyle Marchuk, Matthew F. Krummel

## Abstract

During immune surveillance, CD8 T cells scan the surface of antigen presenting cells using dynamic microvillar palpation and movements as well as by having their receptors pre-concentrated into patches. Here, we use real-time lattice light sheet microscopy to demonstrate the independence of microvillar and membrane receptor patch scanning. While T cell receptor (TCR) patches can distribute to microvilli, they do so stochastically and not preferentially as for other receptors such as CD62L. The distinctness of TCR patch movement from microvillar movement extends to many other receptors that form patches that also scan independently of the TCR. An exception to this is the CD8 co-receptor which largely co-migrates in patches that overlap with or are closely adjacent to those containing TCRs. Microvilli that assemble into a synapse contain various arrays of the engaged patches, notably of TCRs and the inhibitory receptor PD-1, creating a pastiche of occupancies that vary from microvillar contact to contact. In summary, this work demonstrates that localization of receptor patches within the membrane and on microvillar projections is stochastic prior to antigen detection and that such stochastic variation may play into the generation of many individually-composed receptor patch compositions at a single synapse.

**Significance statement:** Motile T cell microvilli palpate surfaces to facilitate surface scanning in a pattern that is independent of the movement of pre-formed patches of transmembrane antigen-receptors across those microvilli; once T cell receptors engage, the microvilli act to scaffold multiple receptors within a microvillar close-contact.

## Introduction

As a key component of adaptive immunity, T cells continuously survey the surfaces of antigen presenting cells (APCs) through close membrane contact to detect agonist antigens with high sensitivity. T cell signaling is initiated when the T cell receptor (TCR) complex recognizes its cognate peptide bound to the major histocompatibility complex (pMHC) (1, 2). During T cell activation, a T cell undergoes profound spatial re-organization of its membrane, surface receptors and downstream signaling molecules to form the immunological synapse (IS), a corrugated membrane apposition to the APC which is critical for T cell signaling and functioning (3, 4).

Given the time constraint for T cells to interact with APCs - the whole cell contact half-life in vivo is about one minute (5) - they are highly efficient in antigen detection. Recent evidence suggests that T cell membrane microvilli facilitate T cell antigen search by participating in serial and parallel palpation events that each have a half-life on the order of just a few seconds (6). Upon pMHC recognition, TCR-enriched microvilli (6), visible as ‘microclusters’ (7, 8), are ‘stabilized’ with half-lives that are 2-5x that of unengaged ‘scanning’ microvilli (6). Microvillar tips are presumed to facilitate TCR-pMHC binding both because they approach APCs to within 15 nm (6), which is the approximate transmembrane-transmembrane dimensions of TCR-pMHC complexes (9, 10), and because TCRs are observed to accumulate in these zones of close contact in response to normal ligand recognition (6, 11).

Biochemical studies demonstrate that TCR half-life for pMHC is a determinant of initiation of TCR signaling; agonism and profound signaling typically requires TCR-pMHC half-lives exceeding 4-6 seconds (12). Likewise, initiation of TCR-mediated signaling events occur within seconds of TCR interaction with cognate antigen (13). TCRs appear to reside in pre-assembled patches or ‘protein islands’, into which other molecules can enter with timescales on the order of seconds, following receptor engagement (14).

These rapid timescales raise the question of how movements of the surface topology (seconds to minutes) and molecular scanning (seconds) are linked, and how and when these are paired during detection and subsequent microvillar stabilization. Further, beyond the TCR, additional membrane receptor-ligand pairs play important roles in enhancing T cell activation and signaling. This includes co-receptor CD8 or CD4 which also binds to MHCs, and both concentrates them (15, 16) and recruits activating lck kinases to the TCR complex (17). Adhesion receptor binding, such as LFA-1 to ICAM-1, initiates cell-cell adhesion (18), and later concentrates in the peripheral-supramolecular activation cluster (pSMAC) of the IS after T cell activation (3). CD28 binds to B7 molecules and generates a critical signal for the activation of naïve T cells whereas the T cell surface receptor PD-1 is a major co-inhibitory molecule (19).

Evidence shows that most T cell surface receptors, like the TCR, can also exist in pre-organized nanoclusters or protein islands before T cell engagement with pMHC (14). However, most of these previous studies were restricted to either a two-dimensional surface or a thin layer of the T cell, lacking the full three-dimensional view of the entire T cell membrane. Therefore, it is also unclear how various receptors are distributed prior to T cell activation, and how they behave in comparison to the microvillar structures that bring T cells into close apposition where ligands can bind.

In this work, we used lattice light sheet microscopy (20) (LLSM) to visualize TCR distribution and membrane topology on entire effector CD8 T cells in 3D in real time. This method provided the ability to track both microvilli and receptors and notably receptor patches at high spatial and temporal resolution. We complemented this with a supported bilayer-based method, called synaptic contact mapping (6) (SCM), in which quantum dots whose diameter is bigger than that of TCR-pMHC complexes are seeded into bilayers containing TCR ligands to enable likewise visualization of the tips of microvilli along with TIRF level measurement of the receptor patches that occupy those contacts. The results demonstrate the independence of membrane topological scanning and receptor patch scanning and emphasize the heterogeneity of the many isolated signaling patches that, in T cells, are arrayed into distinct spatial regions.

## Results

### T cell receptors form high-density patches and distribute independently from microvilli

To establish a baseline for simultaneous studies of receptor patchiness, patch behavior and microvillar behavior in 3D in real-time, we applied lattice light-sheet microscopy (LLSM) to mouse OT-I T cells. These were surface labeled with both non-stimulatory antibodies to the highly abundant surface molecule CD45 and non-stimulatory antibodies to the TCRαβ subunits. Full-cell data was assembled by collecting 150 scans of the light sheet at 175 nm z-intervals and 0.21 Hz frame-rates. Although we cannot resolve details below ~0.2 μm, LLS proved superior to confocal microscopy (**Fig. S1**) even with deconvolution methods, while those methods were orders of magnitude slower for collection of cell volumes. We also reconfirmed (6) that these detection reagents did not induce or enlarge clusters, by performing identical staining on fixed cells (**Fig. S2**) with indistinguishable results.

With the diffraction-limited resolution of LLSM, both microvilli and local foci of higher TCR density, namely patches, were visualized (**Fig. 1A**). Our results confirmed earlier studies that demonstrate TCRs are non-homogenously distributed into high-density patches on the membranes of effector T cells. Analyzing TCR patches relative to the CD45-highlighted membrane topology, we observed that some TCR patches localize to tips of microvilli (**Fig. 1B**); some TCR patches localize to flatter membrane regions (**Fig. 1C**); and some microvilli occur with no TCR patches nearby (**Fig. 1D**).

**Figure. 1.**
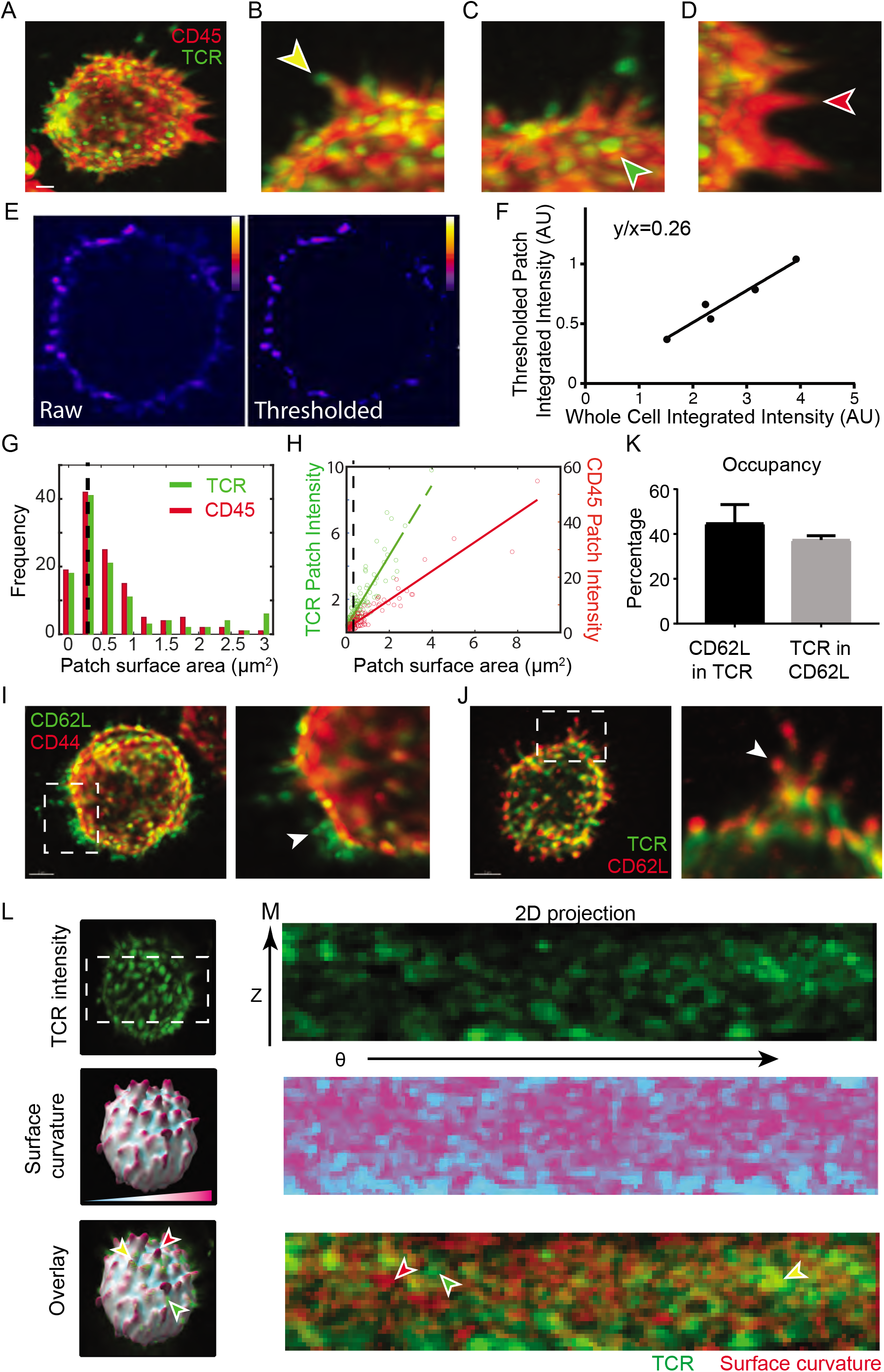
T cell receptors form high-density patches on previously activated T cells and distribute independently from microvilli. (**A)** LLSM image of a previously activated OT-I T cell stained with fluorescent labeled antibodies against CD45 (anti-CD45-Alexa647) and TCRαβ subunits (H57-Alexa488). The surface staining of TCR and CD45 are rendered in green and red respectively. Scale bar: 2 μm. (**B-D)** Distribution of TCR patches relative to microvilli. Zoomed-in regions show examples of (B) TCR patches localized to the tip of a microvillus denoted by the yellow arrow, (**C)** a TCR patch localized outside of microvilli denoted by the green arrow, and **(D)** a microvillus with no TCR patches denoted by the red arrow. **(E)** Slice view of raw (left) and thresholded (right) TCR fluorescence intensity. Thresholded data was used to create 3D surfaces and TCR fluorescence intensity was summed within surfaces to define the thresholded patch integrated intensity **(F)**. **(G)** Size distribution of patches in LLSM. The histogram of patch surface area is obtained from a fixed OT-I T cell stained with fluorescent labeled antibodies against CD45 and TCRαβ subunits. Patch size was determined by using small triangles to cover the edge of the T cell in 3D and calculating the sum of the areas of small triangles that constitute the patch. TCR patch sizes are shown in green and CD45 patch sizes are shown in red. The black dotted line shows the patch size at the diffraction limit. **(H)** Patch surface area and patch fluorescence intensity for TCR and CD45 patches are shown in green dots and red dots respectively. Solid lines are the linear fitted lines for the data. The black dotted line shows the patch size at the diffraction limit. **(I)** Accumulation of CD62L in the microvilli. Left: LLSM image of OTI T cell stained with anti-CD62L-AlexaA488 (green) and anti-CD44-Alexa647 (red). Right: zoomed-in image of the cell region in white dotted box. White arrow points to a microvillus enriched with CD62L. Scale bar: 2 μm. **(J)** TCR patches localize independent of microvilli. Left: LLSM image of an OT-I T cell stained anti-TCRαβ (H57-Alexa488) and anti-CD62L-Alexa647. Right: the zoomed-in image of the cell region in white dotted box. White arrow points to a microvillus enriched with CD62L but not TCR. Scale bar: 2 μm. **(K)** Relationship of TCR and CD62L patches. Fixed OT-I T cells were stained with anti-TCRαβ (H57-Alexa488) and anti-CD62L-Alexa647. Localizations of 266 CD62L patches over 3 cells were used to identify microvilli and measure co-localization of 121 TCR patches using a homebuilt algorithm described in the method section. Black bar: percentage of CD62L patches that overlapped with TCR patches. Grey bar: percentage of TCR patches overlayed with CD62L patches. **(L)** Comparison of TCR patch localization in relation to membrane curvature. OT-I T cells were fixed and then stained with anti-TCRαβ (H57-Alexa488) and anti-CD45-Alexa647. Top to bottom: TCR (green), surface curvature obtained from CD45 intensity data (color map), overlay of TCR and surface curvature. **(M)** 2D Projections of cell volumes from (J). For ease of visualization, overlay is shown with TCR in green and curvature in red. Arrows denote sites where TCR patches locate on microvilli (yellow), on flatter membrane regions (green) and where microvilli contain no obvious TCR patches (red).

Not all TCRs are in patches. This could best be visualized with image thresholding (**Fig 1E**). Applying surfaces to regions above a consistent intensity threshold, we sought to determine what fraction of the TCR was involved in such patches. We thus compared the fraction of receptors in these patches to the total intensity of TCR, taking advantage of the very high signal to noise ratio of LLS. This demonstrated that, across a range of cellular staining intensities, approximately 26% of TCRs are involved in high-density patches (**Fig 1F**). We do not believe that these accumulations were due to membrane folding as they were not coincident with higher distributions for other receptors such as CD45 (**Figure 1A-D**) nor others (see also Figure 3).

**Figure 2.**
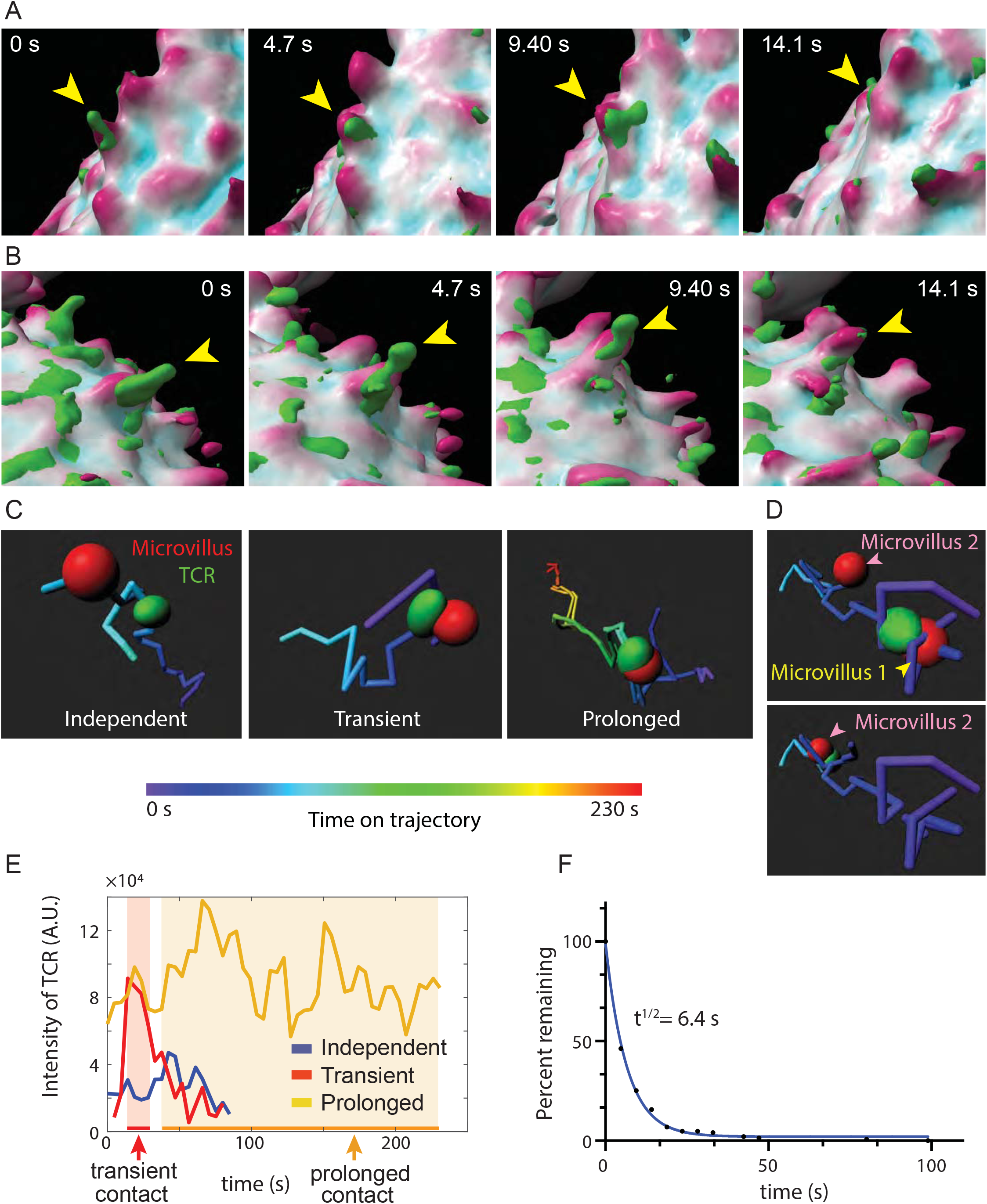
TCR high-density patches move independently relative to microvilli. **(A)** Time series of a TCR patch (yellow arrow) moving over a microvillus. The T cell surface is rendered based on fluorescent signals of CD45. TCR patches labeled with anti-TCRαβ are shown in green. The surface is color-coded to show the curvature, with cyan showing the concave regions and magenta showing the protrusions. **(B)** Time series of a TCR patch (yellow arrow) transiently interacting with a microvillus during the time course of 14 seconds. **(C)** Example trajectories of TCR patches that are independently (left), transiently (middle), or prolonged (right) associated with microvilli. The trajectories are color-coded according to time. Data are extracted from the LLSM movie of a live OT-I T cell stained with anti-CD45-A647 and anti-TCRαβ-A488. TCR patches and microvilli are tracked using the particle tracking function in Imaris. **(D)** The trajectory of a TCR patch that sequentially interacts with two distinct microvilli. **(E)** Fluorescence intensity of TCR patches during interaction with microvilli. Blue: The fluorescence intensity of a TCR patch that is independent of microvilli. Red: The fluorescence intensity of a TCR patch that is transiently associated with a microvillus. Yellow: The fluorescent intensity of a TCR patch that is in prolonged associated with a microvillus. **(F)** Decay graph of the association microvilli with TCR patches. Graph represents interactions taken from a single cell and representative of 3 individual cell measurements.

**Figure 3.**
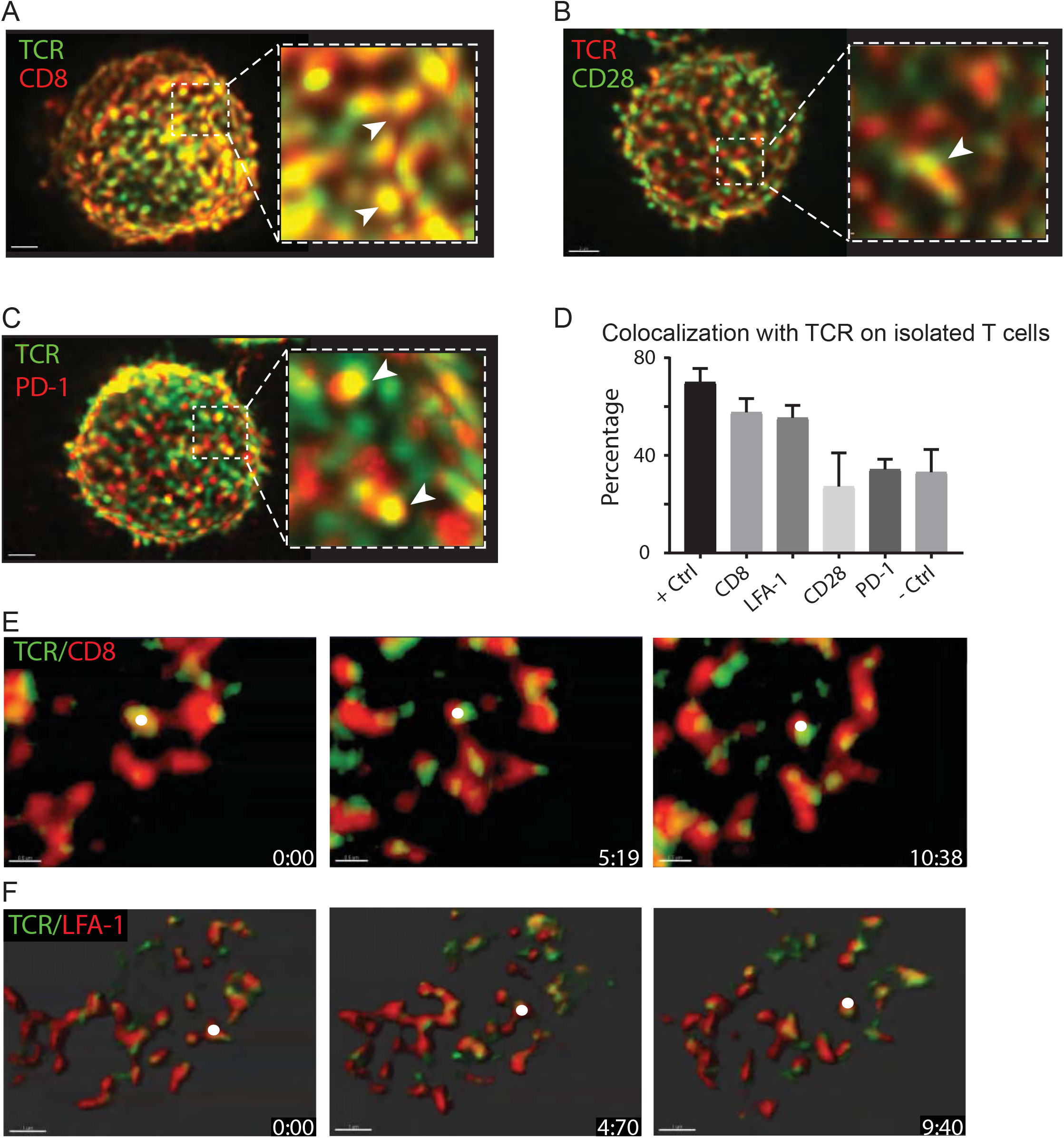
Independent patch-like distributions of surface receptors relative to TCRs. **(A)** The LLSM image of a fixed OT-I T cell stained with anti-TCR-Alexa488 and anti-CD8-Alexa647. The image shows the front half of the cell to demonstrate the overlap between TCR (green) and CD8 (red). The region in the white box is zoomed-in to show more details, and the white arrows mark examples of colocalization of CD8 patches (red) with TCR patches (green). Scale bar: 2 μm. **(B)** The LLSM image of a fixed OT-I T cell stained with anti-TCR-Alexa488 and anti-CD28-Alexa647. The image shows the front half of the cell to demonstrate the overlap between TCR (red) and CD28 (green). The region in the white box is zoomed-in to show more details, and the white arrow marks an example of colocalization of CD28 patches with TCR patches. Scale bar: 2 μm. **(C)** The LLSM image of a fixed OT-I T cell stained with anti-TCR-Alexa488 and anti-PD-1-Alexa647. The image shows the front half of the cell to demonstrate the overlap between TCR (green) and PD-1 (red). The region in the white box is zoomed-in to show more details, and the white arrows marks examples of colocalization of PD-1 patches (red) with TCR patches (green). Scale bar: 2 μm. **(D)** Percentage of patches colocalizing with TCR patches on isolated T cells for each receptor. Receptor patches are segmented from the surface of the cells and a co-receptor patch with over 20% surface area overlap with a TCR patch is considered as colocalized with the TCR patch. OT-I T cells stained with different clones of antibodies to TCR (H57-A488 and TCR-vα-Alexa647) are used as positive controls. OT-I T cells stained with antibodies to CD44 and CD62L are used as negative controls. **(E)** The time series of LLSM images of a live OT-I T cell stained with anti-TCR-Alexa488 and anti-CD8-Alexa647. TCRs are shown in green and CD8a patches are shown in red. The white dot denotes an example of a CD8 patch that colocalize and move with a TCR patch. Scale bar: 0.5 μm. **(F)** The time series of LLSM images of a live OT-I T cell stained with anti-TCR-Alexa488 and anti-CD11a-Alexa647. TCRs are shown in green and LFA-1 patches are shown in red. The white dot denotes an example of an LFA-1 patch that colocalize and move with a TCR patch. Scale bar: 1 μm.

TCR distribution in patches was also observed on a human T cell line (Jurkat) and on naïve OT-I T cells (Fig S2). The latter notably lack detectable microvilli and yet had patchy TCR distributions.

When we measured the size of TCR patches (**Fig. S3A**) as resolved by LLS, we found that their surface area ranged from below diffraction limit (**Fig. S3B**) up to 3 μm^2^ (Fig 1E). CD45, though denser overall, also had patch-like fluctuations of local intensity and these had similar size-distributions (**Fig 1G**). Further, the intensities of fluorescent signals within both TCR and CD45 patches increased proportionally with increasing patch surface area, suggesting a uniform molecular density within patches (**Fig. 1H**).

As a control for microvillar occupancy by a receptor, we similarly labeled CD62L, a molecule which has previously been found to exclusively localize to microvilli (21). Unlike TCR, CD62L distributions were predominantly on the tips of protrusions (**Fig. 1I-J**). Comparing colocalization of TCR and CD62L patches, we found that 45% of CD62L patches co-localized with TCR patches (**Fig. 1K**). Viewed reciprocally, only 38% of TCR patches were found to colocalize with CD62L patches.

To visualize the distribution of TCR patches and microvilli curvature across the entire cell surface, we projected the 3D cell surfaces for each (**Fig 1L**) into 2D images using a radial transformation (**Fig. 1M**). The intense CD45 signal allowed for mapping of surface curvature, and this could be viewed as a color-coded image (Surface Curvature) or overlaid with TCR intensity as a two-color projection (Overlay). Examination of overlaid curvature and TCR projections demonstrated again that only a small fraction of TCR patches overlap with high curvature regions (microvilli). Similarly, when we calculated local surface curvature at locations of TCR patches, we found only a slight preference for those patches to localize to convex surface regions (Fig. S3C). Quantifying TCR patch occupancy by CD45 surface curvature gave ~1/3 occupancy of microvillar tips with TCR patches (**Fig. S3D**), similar to what was quantified using CD62L to mark microvillar tips.

### TCR high-density patches move independently relative to microvilli

A key question was whether these patches were stable or might just be stochastic accumulations, and if they were stable, how would they behave relative to motile microvilli. We thus imaged membrane deformation and TCR on live OT-I T cells with LLSM at frame rates of 0.21 Hz. We found that TCR distributions and microvilli were both highly dynamic (**Movie 1**) and when we tracked the movement of TCR patches and nearby microvilli, we found that TCR patches moved independently from microvilli (**Fig. 2A, B** in which TCR ‘falls off’ a moving microvillus, **Movie 2A**).

When tracking trajectories of microvilli and TCR patches, we found that we could define TCR patches that moved independently of microvilli, transiently associated with microvilli, and in rare cases were in prolonged association with a microvillus (**Fig. 2C and 2D, Movie 2B-D**). We frequently observed cases where TCR patches could be tracked as they moved between microvilli (**Fig. 2D**). Examples of TCR intensities at microvillar tips for each of these forms of interaction were plotted in **Fig. 2E.** Using the tracking data, we calculated the lifetime of association of these patches with microvilli and found an off-rate (t^1/2^) of about 6s (**Fig. 2F**), about the amount of time that a microvillus in general dwells in any one region of space. We did not observe coordinated directionality of TCR patch movement relative to each other. These results show that TCR patches dynamically circulate in and out of microvilli during surface scanning, and it is tempting to imagine the membrane as a sort of skin through which some receptors may circulate, as the cytoskeletal ultrastructure moves beneath.

### Independent dynamic patch-like distributions of co-receptors relative to TCRs

Using this method, we next sought to compare how other membrane receptors were organized relative to microvilli and TCR patches. To determine the overall distribution and avoid using antibodies under conditions that might alter membrane behaviors, we fixed OT-I T cells and labeled them with fluorescent antibodies to CD8, CD28, PD-1 and LFA-1 (CD11a), in combinations with antibodies to TCR, akin to what we had done with CD62L.

CD8, the co-receptor, was found in patches, and of the proteins analyzed in this study, was most frequently found near or co-incident with TCR patches (**Fig. 3A,D**). In contrast, CD28 and PD-1 also formed clusters but neither preferentially co-localized with TCR patches in the steady state (**Fig. 3B-D**). To verify our experimental and analysis procedure for colocalization, we included a positive control experiment in which T cells were labeled with two different clones of antibodies to TCR, providing a measure of the high end of colocalization (**Fig. 3D**). Comparing across the receptors analyzed, LFA-1 segregated with a propensity to co-localize with the TCR, to a similar extent as CD8 (**Fig. 3D**, S4B). While modestly unexpected because the dimensions of LFA-1-ICAM are longer than those of the TCR, such localization may put LFA-1 in a position to quickly participate in adhesion of a microvillus tip while TCR serially engages antigens.

With respect to co-receptor CD8 patches, which were frequently found near TCR patches, we then performed live-imaging at ~0.21Hz, tracking both CD8 and TCR patches with LLSM. This showed that CD8a patches typically were either overlapping or ‘followed’ the movement of TCR patches on isolated OT-I T cells before antigen engagement (**Fig. 3E** and **Movie 3).** Similar prolonged association was observed for TCR with LFA-1 (**Fig. 3F** and **Movie 4**).

### Occupancy and Convergence of TCRs and PD-1 to microvilli tips upon antigen detection

To study the co-assembly of TCR and microvilli location after antigen recognition, we first used Synaptic Contact Mapping (SCM)^7^ whose temporal resolution of the contact region is superior to LLS since all the key data is taken in a single plane. As previously, we incorporated fluorescent Qdots with radii of approximately 16nm into a supported lipid bilayer (SLB) bearing ligands for T cells (**Fig. 4A**). When T cells made close contact with the bilayer, these fluorescent Qdots were sterically excluded from the contact region leaving a dark microvillar “foot-print” of the contact region (**Fig. 4A-B,G**). This allowed us to track both TCR patches and the nearest microvilli contacts on lipid bilayers at time resolution between 1-2 seconds.

**Figure 4.**
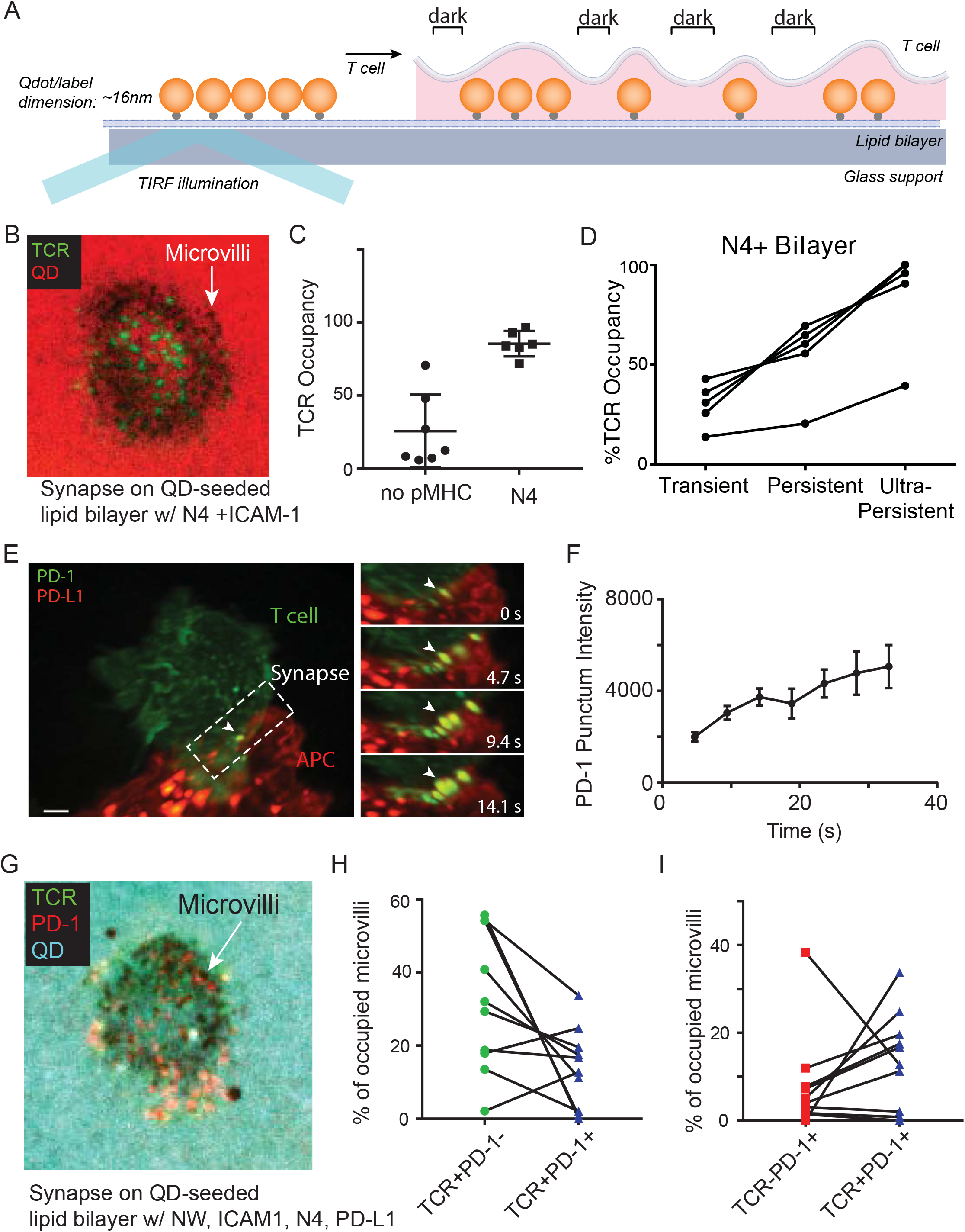
TCR patches and PD-1 patches converge to microvilli tips upon antigen detection. **(A)** Principle of Synaptic Contact Mapping (SCM). Fluorescent Qdot beads of approximately 16nm diameter are incorporated into lipid bilayers which are then seeded with ligands for T cells. Qdot exclusion from a region occurs when a microvillus approaches the bilayer leading to exclusion of Qdots from this region and a ‘hole’ in the otherwise flat fluorescent field formed by the Qdots. **(B)** SCM image of a synapse of an OT-I T cell formed on lipid bilayer containing ICAM1 and N4. OT-I T cells are stained with anti-TCR-Alexa488. TCR microclusters are in green, Qdot signal is in red. Dark spots (as indicated by white arrow) show the locations of microvilli contacting the bilayer where Qdots are sterically excluded. **(C)** TCR occupancy from SCM images for cells interacting with bilayer containing activating N4 pMHC (85.5% ± 3.5% (mean ± SEM), n=7) and cells interacting with bilayer containing no activating N4 pMHC (25.6% ± 9.4% (mean ± SEM), n=6). **(D)** TCR occupancy of microvilli on activating bilayer with N4 pMHC segmented by transient, persistent and ultra-persistent microvilli. Transient microvilli contacts were defined by contacts with persistence time shorter than 3 sigma above the average persistent time of TCR^-^ contacts (cutoffs ranging from ≤ 4-10 s between cells). Persistent microvilli contacts were defined by contacts with persistence time greater than 3 sigma above the average persistent time of TCR^-^ contacts (cutoffs ranging from ≥ 4-10 s between cells). Ultra-persistent microvilli contacts were defined by contacts with persistence time over 90 s. **(E)** PD-1 accumulation at contacts during synapse formation by LLSM. Image of a Jurkat cell genetically expressing PD-1-neonGreen (green) interacting with a CHO cell expressing PD-L1-mScarlet (red) and TCR activator. The synapse region in white dotted box is zoomed and the time series of the region is shown on the right. Scare bar: 2 μm. **(F)** Increase of fluorescence intensity of PD-1 punctum from (E). PD-1 puncta (n=5) were tracked using Imaris and mean intensity was measured for each punctum over time. **(G)** SCM image of a synapse of an OT-I T cell formed on lipid bilayer containing ICAM1, N4 and PD-L1. TCR signal is shown in green, PD-1 signal is shown in red and Qdot signal is shown in cyan. Dark spots (as indicated by white arrow) show the locations of microvilli contacting the bilayer where Qdots are sterically excluded. **(H)** Percentage of microvilli on synapses (n=9) occupied by TCR clusters only (green) and both TCR and PD-1 clusters (blue). **(I)** Percentage of microvilli on synapses (n=9) occupied by PD-1 clusters only (red) and both TCR and PD-1 clusters (blue).

As previously observed (6), most microvillar contacts did not have profound TCR occupancy in the absence of stimulating ligands as measured here by LLS (Fig. 1). This number rose to 85.5% when bilayers contained activating pMHCs compared to 25.6% when bilayers contained no activating pMHCs (**Fig. 4B-C**) and previous work has shown that this is dependent on the agonist peptide and not simply the MHC (6). Further, when we quantified the stability of close contacts on stimulating bilayers, we were able to detect a difference in TCR occupancy between those that were short-lived (persistence time shorter than 3 sigma above the average persistence time of TCR^-^ contacts), persistent (persistence time greater than 3 sigma above the average persistence time of TCR^-^ contacts), and ultra-persistent (defined here as persistence greater than 90 s). In the same synapse, TCR occupancy in super stable close contacts that had persistence times over 90 s was increased on average 2.9 times compared to transient contacts (**Fig. 4D**).

Finally, we took the example of PD-1 to determine the extent to which receptors of similar dimensions partitioned together in the IS. Imaging this receptor and its ligand in real-time in a fluorescent protein-transfected cell system, now using LLSM imaging, we found that PD-1 patches accumulate at the initial contact sites as puncta (**Fig. 4E** and **Movie 5**) and increase in intensity during synapse formation (**Fig. 4F**), suggesting that more receptors are brought into the same contact, rather than one contact having a fixed amount of receptor.

To observe this in the SCM setting and also track whether TCR patches become inhabited by PD-1, we labeled both receptors on T cells placed onto bilayers containing ICAM-1, activating pMHCs and recombinant mouse PD-L1 (**Fig. 4G**). After 3-5 minutes of interaction, we found that PD-1 clusters were typically found in Qdot low regions indicating that this receptor-ligand pair also accumulates at the close contacts and consistent with the length of PD-1-PDL1 interaction being on the order of 14nm, shorter than the size of our tethered Qdot. Within these contacts, cells typically had slightly more microvilli that were occupied by TCR alone, with no detectable PD-1 co-clustering as compared to those that were co-occupied by both TCR and high densities of PD-1 (**Fig. 4H**). This indicates that there is no obligate co-migration of these receptors and likely that there is stochastic engagement and assembly of each into a microvillus. To that end, some contacts (approximately 20%) were PD-1+ but TCR low with indistinguishable fractions that were TCR+PD-1+ (p value is 0.26). This might again imply stochasticity of PD-1 cluster entry into a microvillar region although more study will be needed to fully understand this. In sum, we observe that microvillar contacts co-fill with multiple receptors during recognition.

## Discussion

In this study we demonstrate that on effector T cells prior to antigen engagement, TCRs are distributed into high-density patches. The patch area varies between below the diffraction limit to about 3 um^2^, with most patches sized at or below the diffraction limit (~0.2 μm^2^). This suggests that most TCR patches are nano-scale clusters. This finding is consistent with the observation of pre-organized TCR nanoclusters or protein islands on non-activated T cells (14). Interestingly, we observed TCR patches on naïve T cells which have almost no microvilli. This indicates that TCR patch formation is independent of membrane topology. Although TCR patches have a slightly higher probability to locate on membrane protrusions, its localization is not restricted to microvilli. This finding is on the face different from the works of Haran and co-workers which used multi-angle TIRF imaging to conclude that TCR and other receptors are pre-assembled on microvilli (22, 23). One explanation for the discrepancy in these studies is that multi-angle TIRF has poor ability to estimate the relative intensities when moving away from a coverslip. As noted, although much of the TCR is in these patches which have little microvillar localization, a significant portion of the TCRs appear distributed throughout the membrane outside of patches and would be easily detected by TIRF methods. A limitation of our work is that we focus upon patches, and it is possible and indeed likely that receptors that are not contained in patches circulate and will also therefore populate microvilli, even when no patches are found there. Our work clearly demonstrates a distinction between receptors such as CD62L and TCR, as well as between TCR patches and other receptor patches, neither of which obligately is associated with microvilli.

In the immune synapse, it is well established that TCRs aggregate as micro-clusters and subsequently migrate to the center of the synapse upon engaging antigen (8). Prior to this work, it was known that PD-1 colocalizes with TCR upon binding to PDL-1 and that PD-1-TCR colocalization is required for PD-1-mediated inhibition of TCR signaling (24). Our study finds similarly, albeit with the provision that there are multitudes of mixed clusters containing more or less of PD-1 or TCR. However, those studies which relied upon TIRF imaging did not have the ability to visualize how co-receptors are distributed relative to TCRs before antigen engagement. Here we show that, in addition to TCRs, other T cell surface receptors are also distributed into high-density patches on effector T cells prior to activation. This confirms that TCR are non-uniformly distributed on T cell membranes.

Our previous work had shown that T cells palpate a bilayer with microvilli that are only sparsely occupied with TCR. 3D real-time visualization validates that TCR patches do not obligately live in microvilli, while a control molecule known to preferentially inhabit microvilli, CD62L, was found there, consistent with older work (21). Notably, the co-receptor CD8 often migrates together with TCR which may specifically enhance its ability to co-engage MHC-I molecules on opposing cells. Other receptors, notably CD28 and PD-1, also assemble as patches but do so typically independently of TCR patches.

Like microvilli, TCR patches are highly dynamic. Their movements are not restricted by microvilli: most TCR patches are only transiently associated with microvilli and frequently move in and out of microvilli. Thus, T cell microvillar dynamics (6) and molecular search are largely independent. This may provide benefits for T cells to be able to use sparse non-clustered TCRs to discover large depots of pMHC while clustered TCRs could be sensitive to engage limited collections of pMHC through serial triggering.

To that end, previous work had demonstrated pre-clustering of LFA-1 (25) and this integrin has long been known to play both an essential role in early T cell synapse formation, binding prominently away from the TCR and subsequently forming an adhesion ring (p-SMAC) while TCRs centralize (3). Our finding that LFA-1 patches frequently colocalize with TCRs provides a needed piece of information that adhesive force from pre-clustered LFA-1 is often available in extremely close proximity to the TCR patches. In this setting, it is tempting to speculate that recognition of pMHC and rapid local high-avidity LFA-1 interactions can be efficiently temporally linked. Because TCR signaling intermediates can upregulate LFA-1 affinity via inside-out signaling (26), those integrins may be activated at very low pMHC densities and then assist to stabilize a microvillar contact to permit further TCR signaling. Although signaling downstream of ZAP70 is not required for microvillar stabilization in settings of abundant pMHC (6), it is possible that inside-out signaling to LFA-1 increases the T cell’s sensitivity in very low antigen settings. Integrins are thought to ultimately be excluded from a close contact and then migrate to the p-SMAC as described.

In summary, this work defines the independence of surface receptor movement and microvillar movement on effector T cells before antigen detection. Future work should be possible to understand how dual topological/receptor scanning can lead to high sensitivities of receptor recognition.

## Methods

### Mice

All mice were housed and bred at the University of California, San Francisco, according to Laboratory Animal Resource Center guidelines. Protocols were approved by the Institutional Animal Care and Use Committee of the University of California.

### Cell culture

OT-I T cells were maintained in RPMI supplemented with 10% fetal bovine serum, 100 U/mL penicillin, 0.1 mg/mL streptomycin, 2 mM L-glutamine, 10 mM HEPES and 50 μM β-mercaptoethanol (complete RPMI). Single cell suspensions were prepared from the lymph nodes and spleens of OT-I TCR transgenic mice. Splenocytes were incubated in complete RPMI with 100 ng/mL of SIINFEKL peptide for 30 minutes at 37 °C. The splenocytes were washed three times and suspended in media. Lymphocytes and splenocytes were then mixed 1:1 at 2×10^6^ total cells/mL, and 30 mL of the cell mix was transferred into a T75 flask. After 48 hours, 30 mL of media with IL-2 was added to the flask (final concentration of IL-2: 10 U/mL). After 72 hours, cells were removed from the flask, spun down and resuspended into fresh media with IL-2 and held in culture for an additional 48-96 hours before use.

Bone marrow dendritic cells (BMDCs) were maintained in IMDM supplemented with 10% fetal bovine serum, 100 U/mL penicillin, 0.1 mg/mL streptomycin, 2 mM L-glutamine, and 50 μM β-mercaptoethanol (complete IMDM). BMDCs were prepared from the bone marrow of C57BL/6 mice. BMDCs were incubated in complete IMDM supplemented with 50 μL/mL GM-CSF for 6-10 days. Fresh IMDM and GM-CSF were replenished every 48 hours and cells were passaged on day 5 and day 9. BMDCs were then treated with 60 ng/mL recombinant murine IL-4 for 2-3 days before planned usage and matured using 1 μg/mL of LPS for 5h to overnight before use.

Jurkat-NFAT-PD1-neonGreen cells and CHO-TCRact-PDL1-scarlet were Eli Lilly and Company and described previously. Briefly, CHO-TCRact-PDL1-scarlet cells were generated by transduction of CHO-TCRact cells (Promega J1191: aAPC/CHO-K1) with virus from pLVX-PDL1-mScarlet lentiviral vector (synthesized by GenScript) and cultured in F12 supplemented with 10% fetal bovine serum, 200 mg/mL hygromycin B, and 500 mg/mL G418. CHO-TCRact-PDL1-mscarlet cells were passaged when the density reached 80%-90% confluency. Jurkat-NFAT-PD1-neonGreen suspension cells were generated by transduction of Jurkat-NFAT cells (Promega CS176401) with virus from LVX-PD1-mNeonGreen lentiviral vector (synthesized by GenScript) and cultured in RPMI1640 supplemented with 10% fetal bovine serum, 200 mg/mL hygromycin B, 500 mg/mL G418, 1 mmol/L sodium pyruvate, and 0.1 mmol/L minimum essential medium non-essential amino acids. Jurkat-NFAT-PD1-neonGreen cells were passaged when the cell suspension density reached 2×10^6^ cells/mL. Polybrene (Sigma-Aldrich; TR-1003) was used for lentiviral transduction according to the manufacturer’s instructions.

### Cell preparation for imaging

To prepare OT-I T cells for imaging, live cells were harvested using Ficoll-paque, washed with complete RPMI, and then held at 37 °C. To stain receptors and membrane of T cells for live cell imaging, 2×10^6^ cells were spined down and resuspended in 100 μl RPMI. The cells were then stained with combinations of monoclonal antibodies conjugated to Alexa Fluor dyes on ice for 30 minutes. Cells were then rinsed once with complete RPMI. In SCM experiments, live cells were imaged in RPMI (without phenol red) supplemented with 2% fetal bovine serum, 100 U/mL penicillin, 0.1 mg/mL streptomycin, 2 mM L-glutamine, 10 mM HEPES and 50 μM β-mercaptoethanol (imaging media). In LLSM experiments, live cells were imaged in RPMI (without phenol red) supplemented with 10% fetal bovine serum, 100 U/mL penicillin, 0.1 mg/mL streptomycin, 2 mM L-glutamine, 10 mM HEPES and 50 μM β-mercaptoethanol (imaging media). For TCR staining, 2.5 μg H57-597 non-blocking monoclonal antibody conjugated to either Alexa Fluor 488 was used for 5×10^6^ cells. For CD45 staining, 2.5 μg of CD45 non-blocking monoclonal antibody (clone 30-F11) conjugated to either Alexa Fluor 488 or Alexa Fluor 647 was used. For CD8 staining, 2.5 μg of CD8a monoclonal antibody (clone 53-6.7) conjugated to Alexa Fluor 488. To image PD-1 on OT-I T cell, cells were activated with Dynabeads mouse T activator CD3/CD28 beads for 48 hours. This method increased surface PD-1 level and made it possible for detection. For PD1 staining, 2.5 μg of PD-1 non-blocking monoclonal antibody (clone RMP1-30) conjugated to Alexa Fluor 647 was used.

In SCM live cell imaging, 2×10^5^ cells were added to the bilayer well. Once cells began interacting with the bilayer, imaging was initiated. To prepare for SCM fixed cell imaging, 2×10^5^ cells were added to the bilayer well. Cells were allowed to interact with the bilayer for 3-5 min at 37 °C. Cells were then fixed with 20 mM HEPES, 0.2 M sucrose, 4% paraformaldehyde (PFA) and 0.01% glutaraldehyde for 10 min at 37 °C and washed with PBS before imaging.

To prepare for LLSM live cell imaging, BMDCs were wash with IMDM supplemented with 10% fetal bovine serum, 100 U/mL penicillin, 0.1 mg/mL streptomycin, 2 mM L-glutamine, and 50 μM β-mercaptoethanol (complete IMDM) and plated on fibronectin coated coverslips 15 min before use. OT-I T cells were harvested using Ficoll-paque, washed with complete RPMI and then held at 37 °C. 2×10^6^ OT-I T cells were stained with 2.5 μg of CD45 non-blocking monoclonal antibody conjugated to either Alexa Fluor 488 or Alexa Fluor 647 on ice for 30 minutes, then rinsed once with complete RPMI. OTI cells are added to coverslips with BMDCs and incubated for 10 min at 37 °C. To prepare for LLSM fixed cell imaging, OTI T cells were harvested using Ficoll-paque, washed with complete RPMI. To reduce the possibility of T cell activation due to antibody crosslinking, T cells were labeled and fixed at 4°C. 2×10^6^ OT-I T cells were stained in cold complete RPMI with different combinations of monoclonal antibodies conjugated to Alexa Fluor dyes on ice for 30 minutes, then washed with cold PBS with 5 mM of EDTA at 4°C for 3 times. Cells are then fixed with 20 mM HEPES, 0.2 M sucrose, 4% paraformaldehyde (PFA) and 0.01% glutaraldehyde on ice for at least 1 hour. Fixed cells were washed with cold PBS and spin down on coverslip coated with CellTak before image. For CD62L staining, 2.5 μg of CD62L monoclonal antibody (clone MEL-14) conjugated to Alexa Fluor 647 was used. For CD11a staining, 2.5 μg of CD11a monoclonal antibody (clone M17/4) conjugated to Alexa Fluor 488 was used. For CD28 staining, 2.5 μg of CD28 monoclonal antibody (clone F28) conjugated to Alexa Fluor 488 was used. Isolated cells were selected for analysis.

### Supported lipid bilayers

Preparation and use of supported lipid bilayers was as previously described(6, 27). Briefly, phospholipid mixtures consisting of 96.5% POPC, 2% DGS-NTA (Ni), 1% Biotinyl-Cap-PE and 0.5% PEG5,000-PE in Chloroform were mixed in a round bottom flask and dried, first under a stream of dry nitrogen, then overnight under vacuum. All phospholipids were products of Avanti Polar Lipids. Crude liposomes were prepared by rehydrating the phospholipid cake at a concentration of 4 mM total phospholipids in PBS for one hour. Small, unilamellar liposomes were then prepared by extrusion through 100 nm Track Etch filter papers (Whatman) using an Avestin LiposoFast Extruder (Avestin). 8-well Nunc Lab-Tek II chambered coverglasses were emersed in 5% Hellmanex at 55°C overnight. The chambers were washed repeatedly with 18 MΩ water. After air dry, 250 ul of 3M NaOH was added to each well and kept at 55°C for 15 min. The chamber was then washed with 18 MΩ water. This was repeated once, then the chamber was washed repeatedly with 18 MΩ water and dried under a stream of compress air. Lipid bilayers were setup on a chambered coverglass by adding 0.25 mL of a 0.4 mM liposome solution to the wells. After 30 minutes, wells were rinsed with 8 mL of PBS by repeated addition of 0.5 mL aliquots, followed by aspiration of 0.5 mL of the overlay leaving approximately 0.25ml. Non-specific binding sites were then blocked with 1% BSA in PBS for 30 minutes. After blocking, 25 ng of unlabeled streptavidin was added to each well and allowed to bind bilayers for 30 minutes. After rinsing, protein mixes containing 63 ng ICAM and 6 ng pMHC in 2% BSA were injected into each well. In experiments where TCR, PD-1 localization was imaged, protein mixes containing 63 ng ICAM, 6.25 ng pMHC and 6.25 ng recombinant mouse PD-L1 with 6-His tag (R&D systems) in 2% BSA were injected into each well in this step. ICAM preparation was described previously while pMHC was provided by the NIH Tetramer Facility. After binding for 30 minutes, wells were rinsed again and 25 ng of QDot-streptavidin. Bilayers were finally rinsed with imaging media before being heated to 37 °C for experiments. Imaging media was prepared by supplementing RPMI with 2% fetal bovine serum, 100 U/mL penicillin, 0.1 mg/mL streptomycin, 2 mM L-glutamine, 10 mM HEPES and 50 μM β-mercaptoethanol (complete RPMI).

### Lattice Light Sheet Microscopy

Lattice light-sheet (LLS) imaging was performed in a manner previously described. Briefly, for imaging living cells, 5 mm round coverslips were cleaned by a plasma cleaner and coated with 2 μg/ml fibronectin and TetraSpec fluorescent beads (0.1μm at 1.8×10^8^ beads/mL) in PBS at room temperature for 1 hour or 4 °C overnight before use. TetraSpec beads were used as fiducial markers for correction of shift between different fluorescent channels. BMDCs were dropped onto the coverslip and incubated at 37 °C, 5 % CO_2_ for 20-30 min. Right before imaging, OTI T cells were dropped onto the coverslip which had BMDCs. The sample was then loaded into the previously conditioned sample bath and secured. For imaging fixed cells, 5 mm round coverslips were cleaned by a plasma cleaner and coated with CellTak with TetraSpec fluorescent beads (0.1μm at 1.8X10^8^ beads/mL) at room temperature for 1 hour then washed with 18 MΩ water and dried. Fixed cells were dropped onto the coated coverslip and spin down at 1400×g for 10 min at 4 °C. The coverslip was then mounted on the microscope for imaging.

Imaging was performed with a 488 nm, 560 nm, or 642 nm laser (MPBC, Canada) dependent upon sample labeling in single or two-color mode. Exposure time was 10 ms per frame leading to a temporal resolution of 2.25 s and 4.5 s in single and two-color mode respectively.

### Confocal Microscopy

Comparison confocal images were taken with an upright Leica Sp8 laser scanning confocal microscope equipped with a white light laser (WLL), acousto-optical beam splitter (AOBS), and 5 detectors including 3 PMTs and 2 Hybrid Detectors (HyDs). The objective was a Leica HC PL APO 63x/1.20 Water immersion objective. The images were collected in sequence between channels. Channel one was excited at 499 nm and collected with a HyD detector gain of 496 and a bandpass of 504-755 nm. Channel two was excited at 653 nm and collected with a HyD detector of 318 and a bandpass of 658-777nm. Images were collected at 200 Hz unidirectional scan speed. The pinhole was optimized at 1 AU for 580 nm emission. Pixel/voxel size was 344 nm (x) 344 nm (y) x 356 nm (z) with no digital zoom and 49 nm (x) x 49 nm (x) x 356 nm (z) for the image with 7x digital zoom.

Deconvolution was performed using the Leica LAS X Lightning Deconvolution module using the ‘Adaptive’ strategy. Actual number of iterations were 2 for no digital zoom and 3 for 7x digital zoom. The refraction index was 1.33 for the water mounting medium.

### Synaptic Contact Mapping (SCM)

The TIRF microscope is based on a Zeiss Axiovert 200M with a manual Laser TIRF I slider. To image nanocontacts and TCRs, image sequences consisting of TCRs (TIRF), QD-streptavidin (widefield), and interference reflection microscopy (IRM, widefield) images were collected. For TIRF images, a 488 nm Obis laser (Coherent) was used for excitation of Alexa Fluor 488-labeled TCRs and a 640 nm laser was used for Alexa Fluor 647-labeled PD-1.

SCM used a combination of TIRF detection of TCR and PD-1 with wide-field excitation/emission of Quantum dots. Widefield QD images were acquired using a 405/10x excitation filter (Chroma Technology) in a DG-4 Xenon light source (Sutter). TCR and QD fluorescence were split using a DV2 with a 565 nm long-pass dichroic and 520/35m and 605/70m emission filters (Photometrics). Split images were collected using an Evolve emCCD in quantitative mode (Photometrics). IRM images were acquired using a 635/20x excitation filter (Chroma Technology), and reflected light was collected onto the long-pass side of the DV2 imaging path. For time lapse image series, image sequences were typically acquired at 1 second intervals.

### SCM based contact segmentation and analysis

Contact detection has been described previously. Briefly, the IRM images were first filtered with a high-frequency emphasis filter, then segmented using an active contour to identify the synapse footprint region on the bilayer. Intensity local minima inside the synapse region with intensities less than the median intensity in the synapse region were detected. Each local minimum was dilated with 3-pixel diameter disk structuring element (a cross) and used as a mask input for active contour segmentation of the Qdot image. If the active contour failed to detect a region in the Qdot image, the contour collapsed and the minima was discarded. After independent segmentation of all regions, regions that shared >50% of their area were merged to produce a final segmentation. Pixel indices for identified contact regions that remained after active contour analysis were saved. Contacts and TCR microclusters were tracked using the Imaris autoregressive motion-tracking algorithm with a 0.25 μm frame-to-frame distance criterion and 1 frame gap allowance.

### TIRF Contact / TCR/PD-1 co-localization

The contact TCR/PD-1 co-location was performed using an in-house Matlab script. Briefly, the contact “footprints” were used as a mask to isolate fluorescence intensity associated with TCRs or PD-1. The intensity in each TCR or PD-1 “object” was then averaged and plotted in a histogram. A Gaussian distribution curve centered at the background fluorescence median was then overlayed. The contacts that fell within 3 sigma of the Gaussian distribution were then considered TCR^-^ or PD-1^-^, while the higher intensity contacts were considered TCR^+^ or PD-1^+^ and the binary representations were split into two separate image stacks.

### Lattice Light-Sheet: Post Processing

#### Deconvolution and channel-shift correction

Raw data were deconvolved utilizing the iterative Richardson-Lucy deconvolution process with a known point spread function that was recorded for each color prior to the experiment. The code for this process was provided by the Betzig lab at Janelia Farms. It was originally written in Matlab (The Mathworks) and ported into CUDA (Nvidia) for parallel processing on the graphics processing unit (GPU, Nvidia GeForce GTX Titan X). A Typical sample area underwent 15-20 iterations of deconvolution.

Channel shift between red and green channels were corrected using 4 color-beads embedded on the coverslip. Regions of interest (ROI) within the sampling area were cropped down to size and compressed to 16-bit TIFFs using in-house Matlab code to allow immigration into Imaris (Bitplane).

#### Cluster detection and segmentation

Detection of clusters in images began by detecting the outer edge of cells were detected using a custom program which applied a two-step kmeans clustering calculation on each image slice. Following the identification of the edge of the cell, the detected edge of the cell was, such that the edge being analyzed had a thickness of 3 pixels. In order to locate the approximate positions of puncta, the exterior of the cell was subjected to a Pearson correlation calculation. In this step, a continuous set of small windowed 2d projections were created and a correlation calculation was conducted comparing these projections to a set of gaussian kernels, each of different size. The correlation coefficients were then used in a custom clustering algorithm that applied a 3d watershed calculation to identify the edge and extent of specific clusters.

#### Patch size (area) calculation

To calculate the surface area of patches, the stack of images and stack of images containing the patches were rendered as surfaces. The patch information was then added to the nodes (instead of intensity) after the initial creation of the surface. In a surface, there are many small triangles that comprise the surface. We then find all of the triangles (surface elements) that comprise a patch. The area of each triangle is half the magnitude of the cross product of the vectors that define the triangle. For a patch, we find the area of the constituent triangles and sum them to define the area of the patch.

To determine the size of a diffraction-limited object, a cropped bead image from LLSM was used and added to the 3D T cell image by projecting axially on the edge of the T cell. When adding the bead on the edge, we made a cluster. We then added the bead to different locations on the T cell surface and calculated the surface area of the resulting clusters (the projected beads). The surface area was calculated using the surface rendering (not intensity stack). Every element of the surface is a series of 3 nodes that make up a triangle. Many of these triangles make up the surface. We found all the triangles associated with the bead (cluster) and computed their surface area. The area of each triangle is defined as the half the magnitude of the cross product of the vectors that define the triangle.

#### Cluster colocalization analysis

Colocalization analysis began by registering the images collected in the green and red channels. In this step, any section of a boundary located in the red channel and green channel which were within a radial distance of 3 pixels were considered to be the same. A voxel-based colocalization calculation was then done which identified the voxels and clustered which overlapped in the green and red channels. Finally, a blob analysis yielded sets of colocalizing clusters occupying similar shapes on the exteriors of the images of cells. All algorithms for identifying clusters and calculations with colocalization were written in MatLab (Natick, MA).

#### Defining fraction of TCR in patches

Region of interest was cropped tightly to the isolated cell area, and total TCR fluorescence intensity of that region was used to define the whole cell integrated intensity. To define the thresholded patch integrated intensity, all TCR intensity values below 120 AU were set to 0 to define a consistent threshold across all samples for defining areas of high fluorescence intensity. Surfaces were then created based on thresholded data and TCR fluorescence intensity within those surfaces was summed to get the thresholded patch integrated intensity.

## Author contributions

E.C. performed the majority of experiments, data analysis, figure generation and the formulation of the manuscript. C.B. and J.E. analyzed data and generated figures. K.M. participated in data collection and interpretation. C.B. wrote and edited the manuscript. M.F.K. framed the study, provided scientific direction, analyzed data and wrote the manuscript.

## Acknowledgements

We thank members of the Krummel lab for discussion and Erron Titus for critical reading. This work was supported by NIH grant (MFK) AI052116.

## Supplementary

**Supplementary Figure 1:**
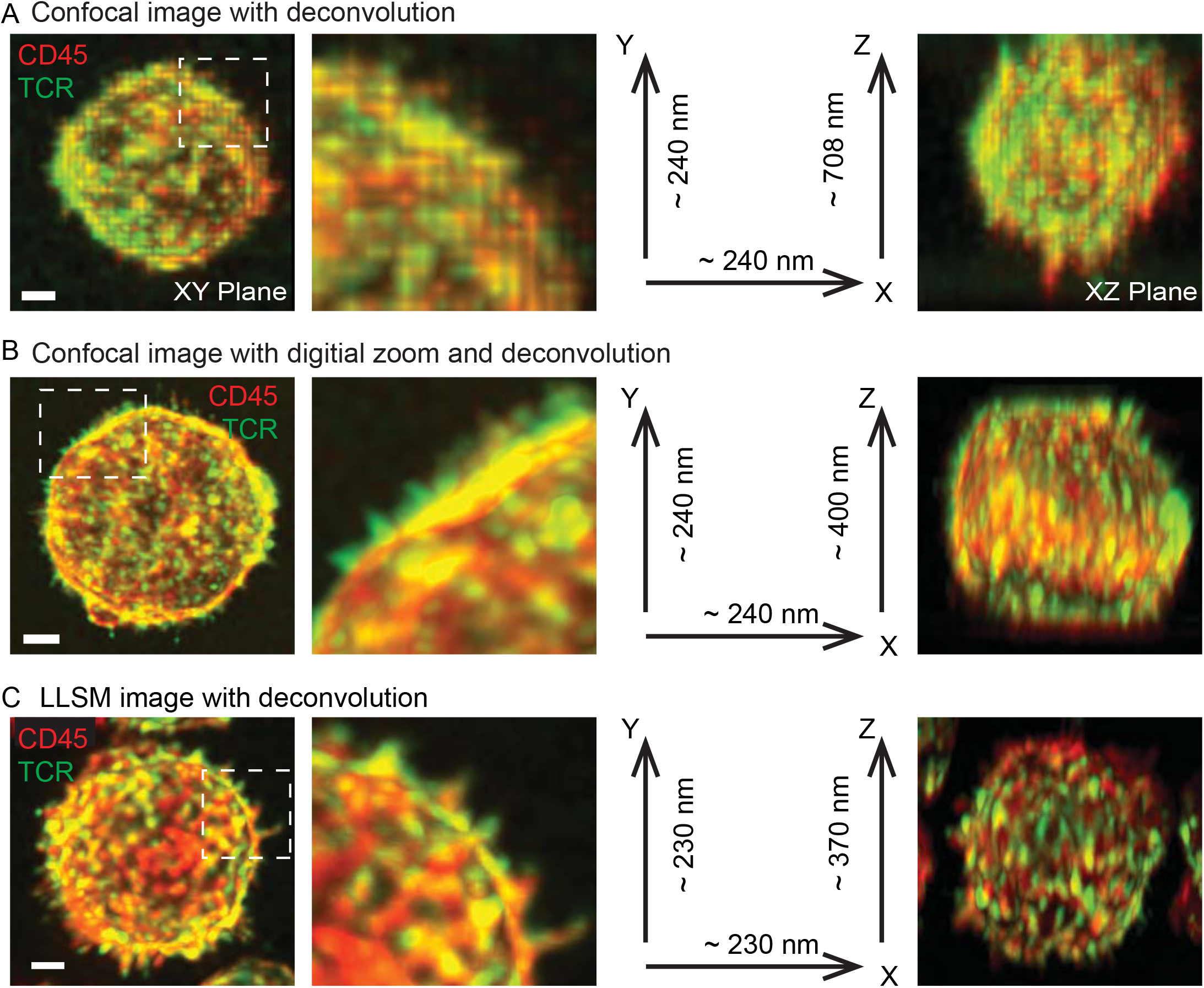
Imaging T cell membrane topology and receptor distributions with confocal microscopy and lattice light sheet microscopy. (A) An OT-I T cell imaged with Nikon SP8 laser scanning confocal microscope at 60X. T cells are stained with anti-CD45-Alexa647 and anti-TCR-Alexa488. Green: TCR signal. Red: CD45 signal. Left: image of a cell in the XY image plane. The boxed region is zoomed in to show the details of the cell. Axes are labeled with estimated resolution. Right: the fluorescence image of the cell in the XZ plane. Scale bar: 2 μm. (B) An OT-I T cell imaged with Nikon SP8 laser scanning confocal microscope with digital zoom. The boxed regions are zoomed in to show the details of the cell. Axes are labeled with estimated resolution. Scale bar: 2 μm. (C) An OT-I T cell imaged with the LLSM at 60X. The boxed regions are zoomed in to show the details of the cell. Axes are labeled with estimated resolution. Scale bar: 2 μm.

**Supplemental Figure S2:**
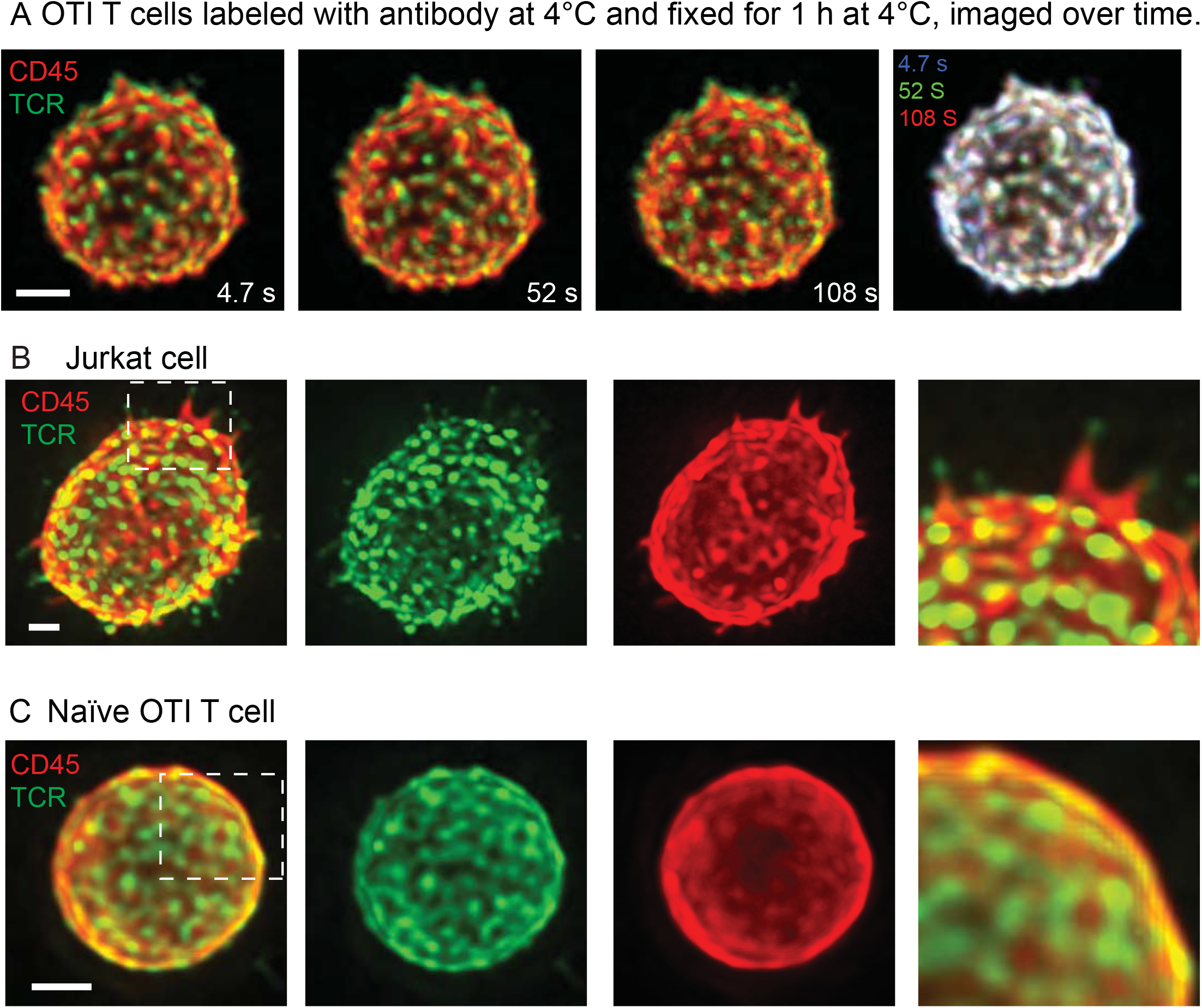
T cell staining and fixation at 4 °C and LLSM images of fixed Jurkat cell and na“ive OTI T cell. (A) Verification of T cell staining and fixation protocol. LLSM image time series of an OT-I T cell stained with anti-CD45-Alexa647 and anti-TCR-Alexa488 at 4 °C and fixed at 4 °C for lh. The time series of the cell has been corrected for photobleaching based on the histogram matching method. Right: overlay of the cells at three time points to show if the TCR patches and microvilli position change over time on fixed cells. The images of the cell at three time points are overlayed after shifting to match for major patterns of the cell. Scale bar: 2 μm. Right: Zoomed in image of the region in box. (B) LLSM image of a fixed Jurkat cell stained with anti-CD45-Alexa647 and anti-TCR-Alexa488. (C) LLSM image of a fixed na“ve OT-I T cell stained with anti-CD45-Alexa647 and anti-TCR-Alexa488. Scale bar: 2 μm. Right: Zoomed in image of the region in box.

**Supplemental Figure S3:**
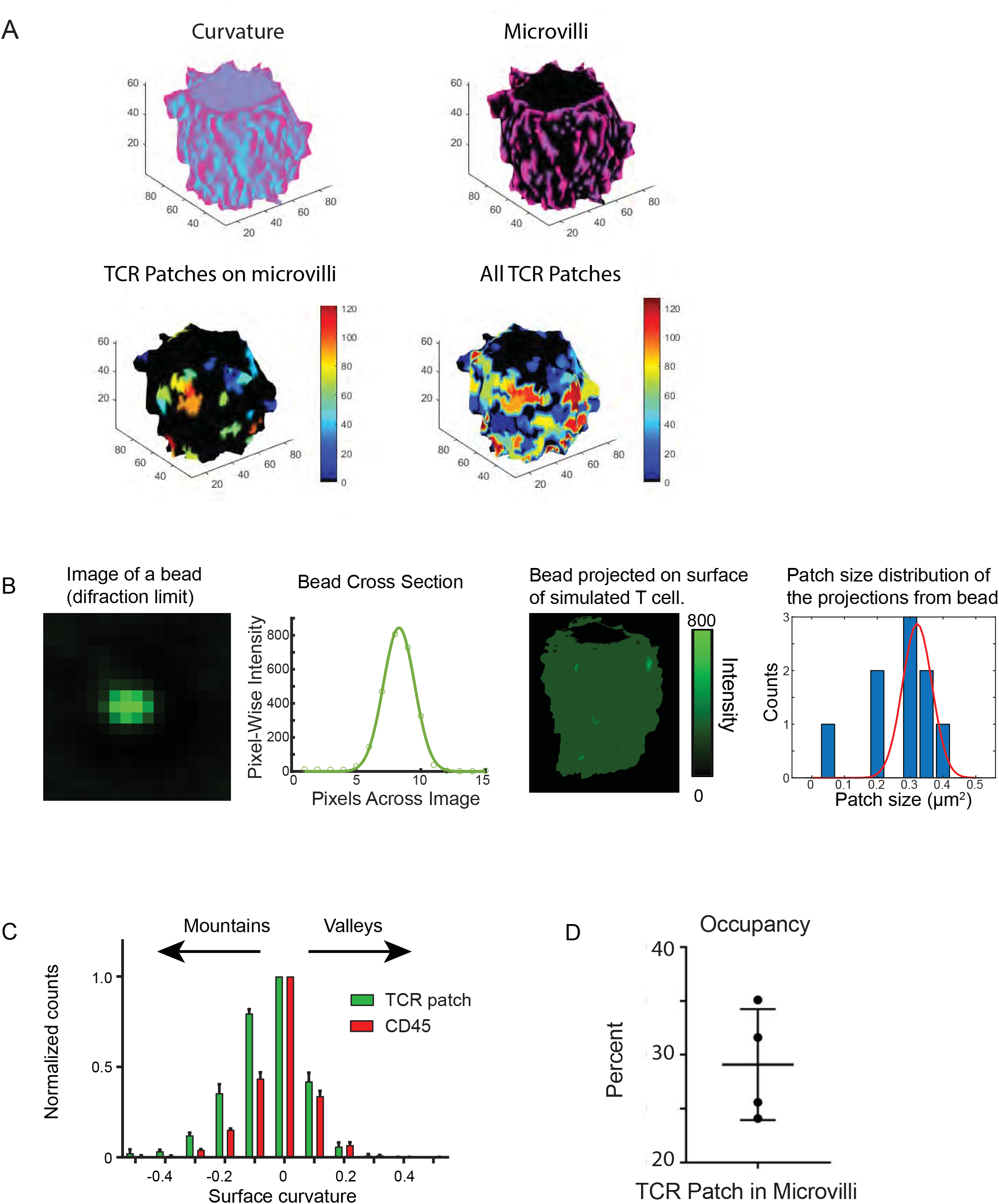
Detection and analysis of microvilli and TCR patches. (A) Microvilli and TCR patch detection on the T cell surface. T cell surface is segmented based on the fluorescence intensity of the surface stained with antibody to CD45. Local curvature is then calculated for the surface with magenta showing convex and cyan showing concave regions. Microvilli are then detected from the local convex regions. TCR patches are segmented based on the fluorescence intensity of surface staining with antibody to TCR. Individual patches are detected and rendered in different colors. TCR patches that have overlap with regions of microvilli can be detected for further analysis. (B) Determination of patch size at the diffraction limit. Left: Image of a diffraction limit sized bead and fluorescence intensity profile of this bead cross section. Right: The projection of the bead on the surface of a simulated T cell and measured patch size of the projection. (C) Distributions of patches. A histogram of TCR and CD45 patches shows their distribution across regions from convex (mountains) to concave (valleys). (D) Occupancy of microvilli by TCR patches. Microvilli are detected based on surface curvature of T cells stained with anti-CD45-Alexa647 as described in (A). TCR patches are detected as described in (A).

**Supplemental Figure S4:**
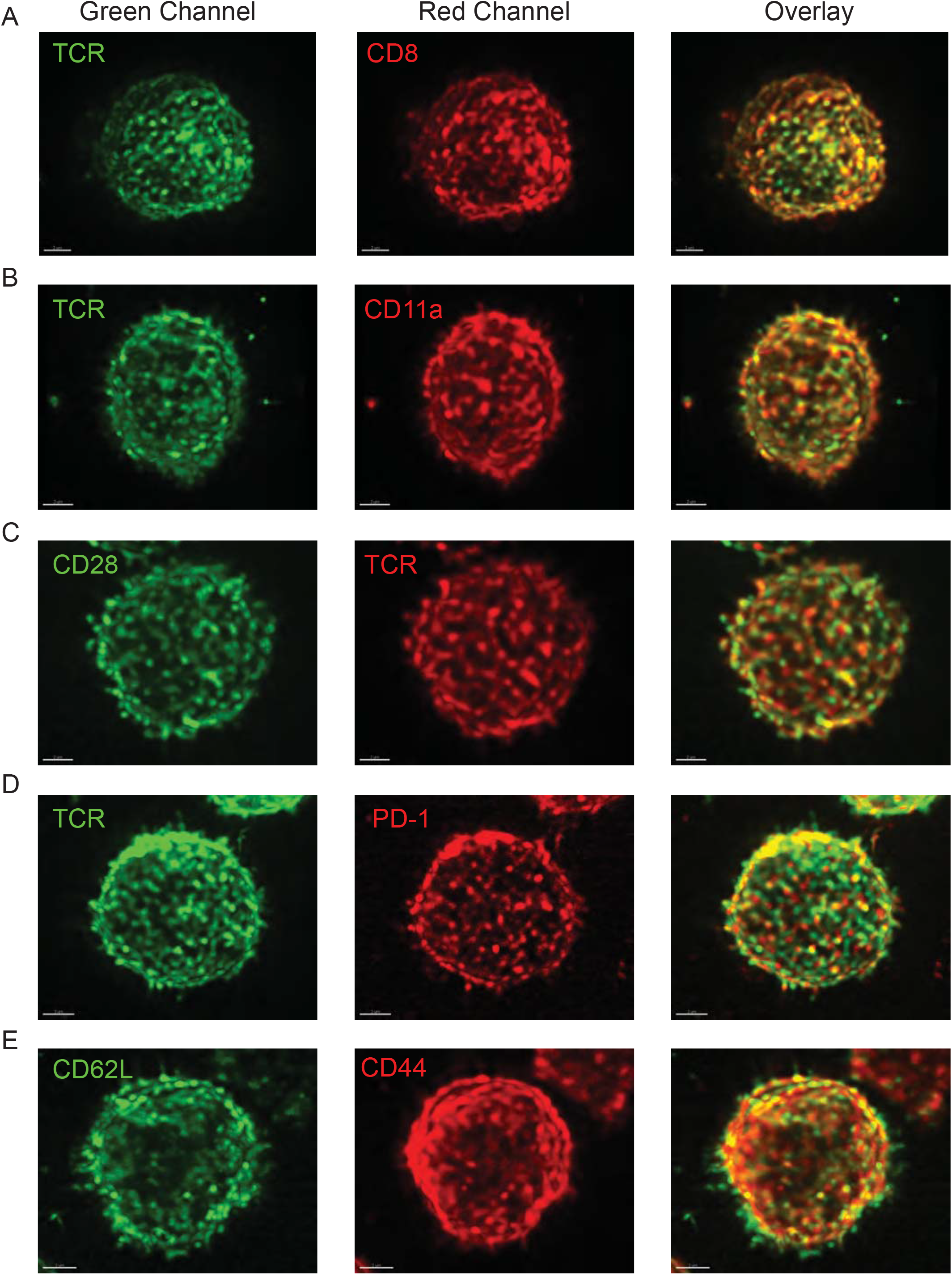
Colocalization of T cell surface receptors with TCR. (A) LLSM image of an OT-I T cell stained with anti-TCR-Alexa488 (H57) and anti-CD8a-Alexa647. Left: TCR channel. Middle: CD8 channel. Right: Overlay of the two channels. (B) LLSM image of an OT-I T cell stained with anti-TCR-Alexa488 (H57) and anti-CD11a-Alexa647. Left: TCR channel. Middle: CD11a channel. Right: Overlay of the two channels. (C) LLSM image of an OT-I T cell stained with anti-TCR-Alexa647 (H57) and anti-CD28-Alexa488. Left: CD28 channel. Middle: TCR channel. Right: Overlay of the two channels. (D) LLSM image of an OT-I T cell stained with anti-TCR-Alexa488 (H57) and anti-PD1-Alexa647. Left: TCR channel. Middle: PD1 channel. Right: Overlay of the two channels. (E) LLSM image of an OT-I T cell stained with anti-CD62L-Alexa488 and anti-CD44-Alexa647. Left: CD62L channel. Middle: CD44 channel. Right: Overlay of the two channels.

**Movie S1: Dynamics of TCR patches on the surface of the T cell.**

LLSM movie of an OT-I T cell stained with anti-CD45-Alexa647 and anti-TCR-Alexa488. The T cell surface is rendered based on CD45 surface staining. TCR patches were detected using a surface algorithm in Imaris and shown in green. The surface is color-coded to show the surface curvature, with cyan showing the concave regions and magenta showing the convex regions. The visualization is rendered in ChimeraX.

**Movie S2: Dynamics of TCR patch relative to microvilli.**

This set of movies demonstrates the dynamics of TCR patches relative to microvilli. TCR patches and microvilli are segmented based on local fluorescence intensity of the TCR staining and CD45 staining respectively. They are tracked over time to generate trajectories.

(A) Surface rendering shows a TCR patch (green) moving on the cell surface (red).

(B) Ball and trajectory shows one TCR patch (green) that has no association with the nearby microvillus (red).

(C) Ball and trajectory shows one TCR patch (green) that is transiently associated with the nearby microvillus (red).

(D) Ball and trajectory shows one TCR patch (green) that has prolonged association with the nearby microvillus (red).

**Movie S3: Comigration of TCR and CDS Receptor Patches.**

This movie shows the dynamics of CD8 (labeled with anti-CD8a-Alexa647) and TCR (labeled with H57-Alexa488) during normal scanning of an OT-I blast.

**Movie S4: Comigration of TCR and LFA-1 Receptor Patches.**

This movie shows the dynamics of LFA-1 (labeled with anti-CD11a-Alexa647} and TCR (labeled with H57-Alexa488} during normal scanning of an OT-I blast.

**Movie S5: PD-1 accumulation at the initial contacts during synapse formation.**

This movie shows the accumulation of PD-1 at initial contacts during synapse formation between a Jurkat cell expressing PD1-neonGreen and a CHO cell expressing TCR-activator and PD-L1-mScarlet. Green: PD1 expressed on the Jurkat cell. Magenta: PD-L1 expressed on the CHO cell.

## References

1. M. M. Davis et al., T cells as a self-referential, sensory organ. Annu Rev Immunol 25, 681–695 (2007).

2. P. A. van der Merwe, O. Dushek, Mechanisms for T cell receptor triggering. Nat Rev Immunol 11, 47–55 (2011).

3. A. Grakoui et al., The immunological synapse: a molecular machine controlling T cell activation. Science 285, 221–227 (1999).

4. D. R. Fooksman et al., Functional anatomy of T cell activation and synapse formation. Annu Rev Immunol 28, 79–105 (2010).

5. A. Gerard et al., Detection of Rare Antigen-Presenting Cells through T Cell-Intrinsic Meandering Motility, Mediated by Myo1g. Cell 158, 492–505 (2014).

6. E. Cai et al., Visualizing dynamic microvillar search and stabilization during ligand detection by T cells. Science 356 (2017).

7. G. Campi, R. Varma, M. L. Dustin, Actin and agonist MHC-peptide complex-dependent T cell receptor microclusters as scaffolds for signaling. J Exp Med 202, 1031–1036 (2005).

8. T. Yokosuka et al., Newly generated T cell receptor microclusters initiate and sustain T cell activation by recruitment of Zap70 and SLP-76. Nat Immunol 6, 117–127 (2005).

9. K. Choudhuri, D. Wiseman, M. H. Brown, K. Gould, P. A. van der Merwe, T-cell receptor triggering is critically dependent on the dimensions of its peptide-MHC ligand. Nature 436, 578–582 (2005).

10. S. J. Davis, P. A. van der Merwe, The kinetic-segregation model: TCR triggering and beyond. Nat Immunol 7, 803–809 (2006).

11. K. Y. Chen et al., Trapping or slowing the diffusion of T cell receptors at close contacts initiates T cell signaling. Proceedings of the National Academy of Sciences of the United States of America 118, e2024250118 (2021).

12. D. S. Lyons et al., A TCR binds to antagonist ligands with lower affinities and faster dissociation rates than to agonists. Immunity 5, 53–61 (1996).

13. I. Stefanova et al., TCR ligand discrimination is enforced by competing ERK positive and SHP-1 negative feedback pathways. Nat Immunol 4, 248–254 (2003).

14. B. F. Lillemeier, J. R. Pfeiffer, Z. Surviladze, B. S. Wilson, M. M. Davis, Plasma membrane-associated proteins are clustered into islands attached to the cytoskeleton. Proc Natl Acad Sci U S A 103, 18992–18997 (2006).

15. C. Wulfing et al., Costimulation and endogenous MHC ligands contribute to T cell recognition. Nat Immunol 3, 42–47 (2002).

16. C. R. Glassman, H. L. Parrish, M. S. Lee, M. S. Kuhns, Reciprocal TCR-CD3 and CD4 Engagement of a Nucleating pMHCII Stabilizes a Functional Receptor Macrocomplex. Cell reports 22, 1263–1275 (2018).

17. J. M. Turner et al., Interaction of the unique N-terminal region of tyrosine kinase p56lck with cytoplasmic domains of CD4 and CD8 is mediated by cysteine motifs. Cell 60, 755–765 (1990).

18. C. Wülfing, M. D. Sjaastad, M. M. Davis, Visualizing the dynamics of T cell activation: Intracellular adhesion molecule 1 migrates rapidly to the T cell/B cell interface and acts to sustain calcium levels. Proceedings of the National Academy of Sciences, USA 95, 6302–6307 (1998).

19. B. T. Fife, J. A. Bluestone, Control of peripheral T-cell tolerance and autoimmunity via the CTLA-4 and PD-1 pathways. Immunological Reviews 224, 166–182 (2008).

20. B. C. Chen et al., Lattice light-sheet microscopy: imaging molecules to embryos at high spatiotemporal resolution. Science 346, 1257998 (2014).

21. U. H. von Andrian, S. R. Hasslen, R. D. Nelson, S. L. Erlandsen, E. C. Butcher, A central role for microvillous receptor presentation in leukocyte adhesion under flow. Cell 82, 989–999 (1995).

22. Y. Jung et al., Three-dimensional localization of T-cell receptors in relation to microvilli using a combination of superresolution microscopies. Proceedings of the National Academy of Sciences 113, E5916–E5924 (2016).

23. S. Ghosh et al., ERM-Dependent Assembly of T Cell Receptor Signaling and Co-stimulatory Molecules on Microvilli prior to Activation. Cell reports 30, 3434–3447.e3436 (2020).

24. T. Yokosuka et al., Programmed cell death 1 forms negative costimulatory microclusters that directly inhibit T cell receptor signaling by recruiting phosphatase SHP2. Journal of Experimental Medicine 209, 1201–1217 (2012).

25. A. Cambi et al., Organization of the integrin LFA-1 in nanoclusters regulates its activity. Mol Biol Cell 17, 4270–4281 (2006).

26. Y. Zhang, H. Wang, Integrin signalling and function in immune cells. Immunology 135, 268–275 (2012).

27. P. Beemiller, J. Jacobelli, M. F. Krummel, Integration of the movement of signaling microclusters with cellular motility in immunological synapses. Nat Immunol ni.2364 [pii]10.1038/ni.2364 (2012).

